# Identification of a transient receptor potential channel that is regulated by phospholipid asymmetry

**DOI:** 10.64898/2026.08.08.743638

**Authors:** Riku Nakanishi, Akira Murakami, Emi Sasaki, Masaki Tsuchiya, Miki Suzuki, Akifumi Shiomi, Kohjiro Nagao, Tomohiko Taguchi, Masato Umeda, Kunitoshi Uchida, Yuji Hara

## Abstract

Phospholipid asymmetry is a hallmark of mammalian cell membranes and reflects the selective distribution of distinct phospholipid species between the two leaflets of the lipid bilayer. Although this asymmetry is tightly maintained, the membrane proteins whose functions depend on it remain largely unknown. To perturb phospholipid asymmetry experimentally, we expressed a constitutively active phospholipid scramblase and thereby identified transient receptor potential melastatin 8 (TRPM8) as an ion channel regulated by this membrane property. Activation of TRPM8 by both *l*-menthol and innocuous cold was markedly suppressed following disruption of phospholipid asymmetry. Likewise, selective depletion of phosphatidylserine (PS), a phospholipid enriched in the cytoplasmic leaflet, using a cytosolically targeted PS decarboxylase attenuated TRPM8 activation, indicating that cytoplasmic PS is required for proper TRPM8 function. Mechanistically, our findings suggest that cytoplasmic PS supports efficient TRPM8 activation by maintaining the biochemical state of the channel. Together, these findings identify TRPM8 as a phospholipid asymmetry-dependent ion channel and establish an experimental strategy for systematically identifying membrane proteins regulated by phospholipid asymmetry. This work provides a foundation for future studies investigating the biological significance of this fundamental membrane property.

## Introduction

The cell membrane is a lipid bilayer composed primarily of phospholipids and cholesterol. Structural diversity among phospholipids arises from differences in their polar head groups and fatty acyl chains, and their composition varies across intracellular membranes (1). Phospholipid composition also differs between the two leaflets of the lipid bilayer, a characteristic referred to as phospholipid asymmetry. In the plasma membrane of mammalian cells, phosphatidylcholine (PC) is enriched in the outer leaflet, whereas phosphatidylethanolamine (PE), phosphatidylserine (PS), and phosphatidylinositol (PI) are predominantly localized to the inner leaflet. This asymmetric distribution is actively maintained by phospholipid transporters, including adenosine triphosphate (ATP)-dependent flippases that translocate phospholipids from the outer to the inner leaflet and ATP-independent scramblases that facilitate bidirectional phospholipid movement (2).

Although phospholipid asymmetry is tightly maintained under physiological conditions, its disruption plays essential roles in biological processes (3,4). During apoptosis initiation, platelet activation, and cell-cell fusion, PS becomes exposed on the outer leaflet of the plasma membrane, where it serves as a signaling molecule required for these specific cellular processes. Despite this, the physiological significance of maintaining phospholipid asymmetry under steady-state conditions remains poorly understood. Notably, comparison of the reported maximal PS transport capacity of erythrocytes with cellular ATP turnover suggests that PS transport could account for a substantial fraction of ATP consumption, underscoring the potential physiological importance of phospholipid asymmetry (5,6). Previously, we demonstrated that disruption of phospholipid asymmetry by genetic deletion of the phospholipid flippase ATP11A impairs myotube morphology through suppression of the mechanosensitive ion channel PIEZO1 (7). We further showed that PIEZO1 promotes the proliferation of muscle stem cells prior to their differentiation into myoblasts (8). Together, these observations suggested that phospholipid asymmetry regulates distinct cellular processes through modulation of specific membrane proteins, leading us to propose the “flip-flop switch” hypothesis.

Membrane lipids have long been recognized as regulators of cellular function through modulation of membrane proteins and membrane-associated proteins (9). We have focused on transient receptor potential (TRP) channels, which transduce extracellular stimuli into intracellular signals (10–13). Numerous TRP channels have been identified as targets of lipid regulation. For example, membrane cholesterol regulates TRPV2 (Vanilloid 2) (14) and TRPC5 (Canonical 5) (15,16), whereas phosphatidylinositol 4,5-bisphosphate (PIP₂) directly regulates TRPV1 (Vanilloid 1) (17), TRPV5 (Vanilloid 5) (18), TRPML1 (Mucolipin 1) (19), and TRPM8 (Melastatin 8) (20). Although PIP₂ is a minor phospholipid localized to the inner leaflet of the plasma membrane, these studies have primarily focused on how specific lipid–protein interactions regulate channel activity rather than on the functional significance of phospholipid asymmetry. Consequently, whether phospholipid asymmetry itself regulates membrane protein function remains largely unexplored, not only for TRP channels but also for membrane proteins and membrane-associated proteins more broadly. This gap in knowledge primarily reflects the absence of an experimental strategy capable of systematically identifying proteins regulated by phospholipid asymmetry.

An important finding from our previous study was that overexpression of a constitutively active scramblase recapitulated the reduction in PIEZO1 activity observed in flippase-deficient cells (7). This observation led us to hypothesize that phospholipid asymmetry could be experimentally perturbed in virtually any cell type, thereby enabling systematic identification of membrane proteins regulated by phospholipid asymmetry. Building on this concept and our longstanding interest in TRP channels, we applied this strategy to the TRP channel family and identified TRPM8 as a phospholipid asymmetry-dependent ion channel.

## Results

**A constitutively active phospholipid scramblase-based screen identifies TRPM8 as a potential target of regulation via phospholipid asymmetry.**

Before initiating the screen, we first reassessed whether expression of the constitutively active phospholipid scramblase transmembrane protein 16F (TMEM16F) efficiently disrupts plasma membrane phospholipid asymmetry. Wild-type (WT) TMEM16F requires an increase in intracellular Ca²⁺ for activation, whereas a constitutively active mutant (CA-TMEM16F) containing a 21-amino-acid insertion within the N-terminal region together with the D430G substitution has been reported to exhibit scramblase activity independently of intracellular Ca²⁺ (Fig. 1A) (21,22). WT-TMEM16F, CA-TMEM16F, or the phospholipid scrambling-deficient R499A mutant (23) were bicistronically expressed together with the fluorescent protein mCherry in HEK293T cells (Fig. S1A). PS, which is normally restricted to the inner leaflet of the plasma membrane, was detected at the cell surface by extracellular application of Annexin-V. Annexin-V staining along the plasma membrane was observed exclusively in cells expressing CA-TMEM16F (Fig. S1A). Consistent with this observation, FLAG immunostaining showed that Annexin-V positivity was confined to cells expressing FLAG-tagged CA-TMEM16F (Fig. 1B). We next screened TRP channels to identify those whose activity was altered by coexpression of CA-TMEM16F. For the initial screen, we selected Ca²⁺-permeable TRP channels with well-characterized chemical agonists (24). Following agonist application (capsaicin, GSK1016790A, allyl isothiocyanate (AITC), and *l*-menthol, respectively), HEK293T cells expressing TRPV1, TRPV4, TRPA1 (Ankyrin 1), or TRPM8 exhibited an increase in the Fura-2 ratio, indicating agonist-induced Ca²⁺ influx. Expression of CA-TMEM16F did not affect agonist-induced Ca²⁺ influx mediated by TRPV1, TRPV4, or TRPA1 (Fig. 1C–E and Fig. S2A–C). By contrast, agonist-induced increases in intracellular Ca²⁺ were significantly reduced in cells coexpressing TRPM8 and CA-TMEM16F following *l*-menthol stimulation (Fig. 1F and Fig. S2D). In addition, CA-TMEM16F suppressed TRPM8-dependent Ca²⁺ influx induced by icilin, a structurally distinct TRPM8 agonist (Fig. S3A, B). To determine whether this effect was specific to TMEM16F, we examined TRPM8 activity after expressing XK-related protein 8 (XKR8) (25), a phospholipid scramblase distinct from TMEM16F. Expression of *Aedes albopictus* XKR (AaXKR), which we previously reported to function as a constitutively active, Ca²⁺-independent phospholipid scramblase (26), likewise reduced *l*-menthol-induced Ca²⁺ influx (Fig. S3C, D). Together, these findings suggest that TRPM8 is a TRP channel positively regulated by phospholipid asymmetry.

**Figure 1.**
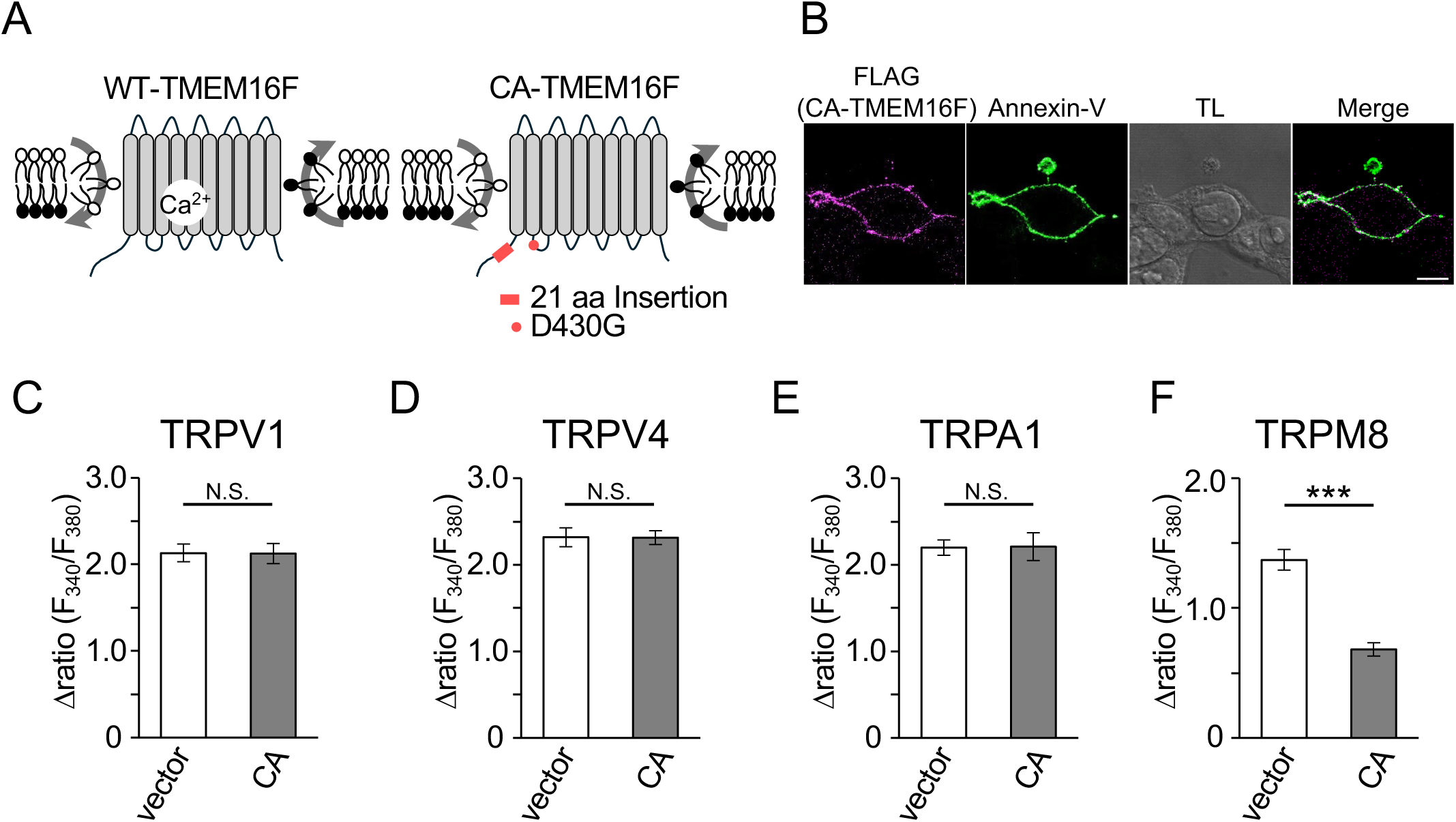
Scramblase-based screen for transient receptor potential (TRP) ion channels regulated by phospholipid asymmetry. (A) Schematic representation of wild-type (WT) and constitutively active (CA) TMEM16F. CA-TMEM16F contains an additional 21-amino-acid sequence within the N-terminal region together with a D430G substitution. (B) Detection of phosphatidylserine (PS) exposure at the plasma membrane in cells expressing CA-TMEM16F. PS exposure was visualized using Annexin V fused to EGFP. (C–F) Agonist-induced Ca²⁺ influx in cells expressing the indicated channels. Ca²⁺ influx was quantified as the maximum Fura-2 Δratio. Mean ± SEM. vector, n = 79, 72, 100, 78 cells; CA, n = 87, 180, 84, 132 cells, respectively. *** *P* < 0.001; N.S.., not significant.

### Agonist-evoked TRPM8 currents require phospholipid asymmetry

To validate the screening results, we performed whole-cell patch-clamp recordings in HEK293T cells expressing rat TRPM8. In control cells, *l*-menthol evoked TRPM8 currents with the characteristic outwardly rectifying I–V relationship (Fig. 2A, F) (27,28). At −60 mV, *l*-menthol-evoked inward currents were significantly smaller in cells coexpressing CA-TMEM16F than in cells expressing TRPM8 alone. Even at a higher concentration of *l*-menthol, CA-TMEM16F suppressed TRPM8 currents (Fig. S4A–C). By contrast, neither WT-TMEM16F nor the scramblase activity-deficient R499A mutant significantly affected *l*-menthol-induced currents (Fig. 2A–F). Together, these results demonstrate that phospholipid asymmetry is required for agonist-evoked TRPM8 activity.

**Figure 2.**
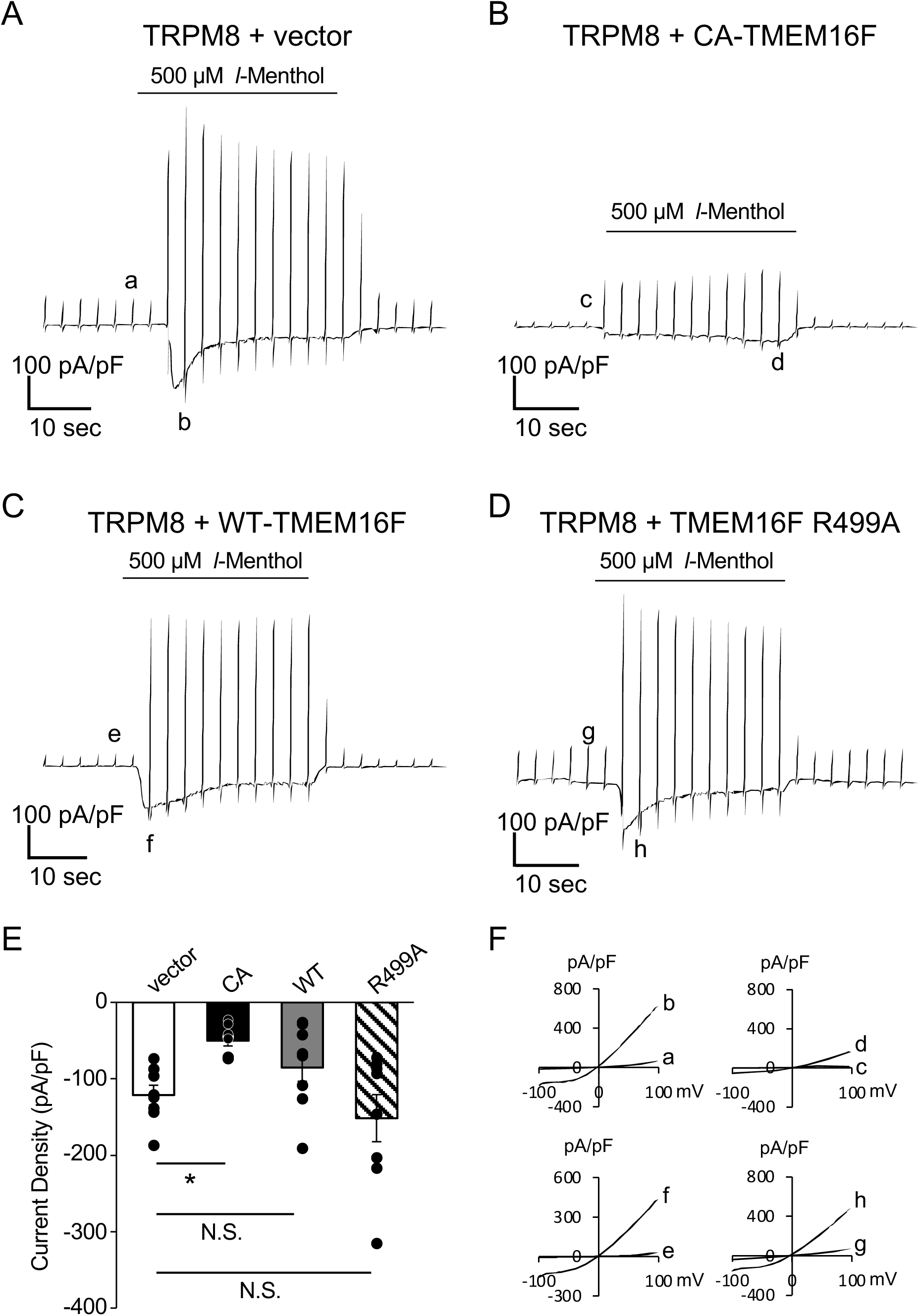
***l*-Menthol-evoked TRPM8 currents in phospholipid asymmetry-disrupted cells.** (A-D) Representative traces of TRPM8 currents evoked by 500 µM *l*-menthol in cells transiently expressing TRPM8 alone (A) or coexpressing CA-TMEM16F (B), WT-TMEM16F (C), or TMEM16F R499A (D). (E) TRPM8 current density induced by *l*-menthol in cells expressing the indicated TMEM16F constructs. Mean ± SEM. (F) I-V relationships obtained from voltage ramp pulses applied at the indicated time points during the recordings (a-h in A-D, respectively). TMEM16F R499A is a scramblase activity-deficient mutant of CA-TMEM16F. \**P* < 0.05; N.S.., not significant.

### Cold-evoked TRPM8 activation requires phospholipid asymmetry

TRPM8 is activated not only by chemical agonist such as *l*-menthol but also by innocuous cold, which has been proposed to gate the channel through a mechanism distinct from that of chemical agonists (28). We therefore investigated whether phospholipid asymmetry is likewise required for cold-evoked TRPM8 activation. As described previously (27), HEK293T cells expressing rat TRPM8 were exposed to cooled solution, and intracellular Ca²⁺ responses and TRPM8 currents were evaluated. Fura-2 Ca²⁺ imaging showed that expression of CA-TMEM16F significantly reduced cooling-induced Ca²⁺ influx (Fig. 3A, B). A comparable reduction was observed for mouse TRPM8, indicating that this effect is not limited to the rat channel (Fig. S5A, B). Consistent with these observations, whole-cell recordings demonstrated that cooling-evoked TRPM8 currents were attenuated by CA-TMEM16F expression (Fig. 3C–G). Together, these findings demonstrate that phospholipid asymmetry is required for TRPM8 activation by both chemical agonists and cold.

**Figure 3.**
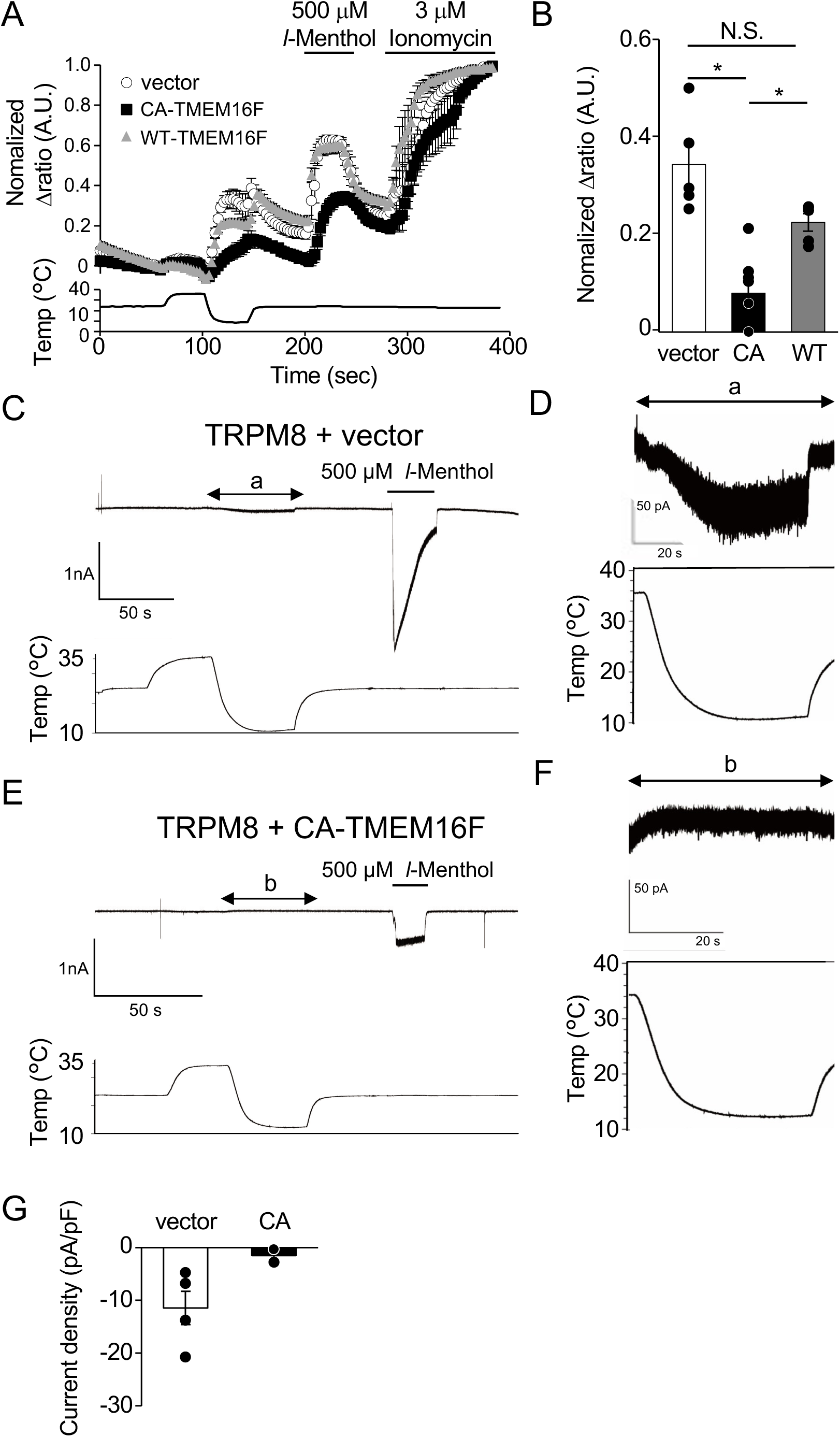
Temperature-dependent TRPM8 activity in phospholipid asymmetry-disrupted cells. (A) Fura-2 ratiometric measurements (F340/F380) of Ca^2+^ mobilization in cells transiently expressing TRPM8 alone (open circles), or coexpressing CA-TMEM16F (filled squares) or WT-TMEM16F (gray triangles). Cells were sequentially exposed to cold stimulation, *l*-menthol, and ionomycin. The Fura-2 ratio was normalized to the value obtained following ionomycin application. Average Δratio traces are shown. The lower panel shows a representative temperature profile recorded adjacent to the cells during Fura-2 imaging. (B) Maximum Fura-2 Δratio during cold stimulation calculated from (A). (C–F) Representative TRPM8 current traces by cold stimulation in cells transiently expressing TRPM8 alone (C, D) or coexpressing CA-TMEM16F (E, F). (G) TRPM8 current density induced by cold stimulation in cells expressing TRPM8 alone or coexpressing CA-TMEM16F (n = 4 and 2 cells, respectively). Mean ± SEM. \**P* < 0.05, \*\*\**P* < 0.001; N.S.., not significant.

### Cytoplasmic leaflet of PS is involved in TRPM8 activation

We next investigated how disruption of phospholipid asymmetry suppresses TRPM8 activity. Because PS is highly enriched in the cytoplasmic leaflet of the plasma membrane but only sparsely distributed in the extracellular leaflet (29), phospholipid scrambling by CA-TMEM16F would be expected to markedly reduce the abundance of PS in the cytoplasmic leaflet. To selectively deplete cytoplasmic PS, we exploited the phosphatidylserine decarboxylase, which converts PS into PE. Although the mammalian phosphatidylserine decarboxylase PISD is predominantly localized to mitochondria, an N-terminally truncated form of the yeast homolog yPSD1 (yPSD1ΔN) has been reported to localize to the cytoplasm while retaining enzymatic activity in mammalian cells (30). We next visualized cytoplasmic PS using the genetically encoded PS probe LactC2-mRuby (31). In control cells, LactC2-mRuby fluorescence was predominantly localized to the plasma membrane. By contrast, this fluorescence signal was significantly reduced in cells expressing yPSD1ΔN but not in cells expressing the catalytically inactive mutant yPSD1ΔN S463A (Fig. S6A, B). These findings indicate that yPSD1ΔN reduces the abundance of PS within the cytoplasmic leaflet of the plasma membrane. We then examined how depletion of cytoplasmic PS affects TRPM8 activity. In TRPM8-expressing cells, coexpression of yPSD1ΔN significantly reduced *l*-menthol-induced Ca²⁺ influx, whereas coexpression of the catalytically inactive mutant yPSD1ΔN S463A produced only a minor effect (Fig. 4A, B). Together, these findings suggest that PS in the cytoplasmic leaflet of the plasma membrane is required for TRPM8 activity.

**Figure 4.**
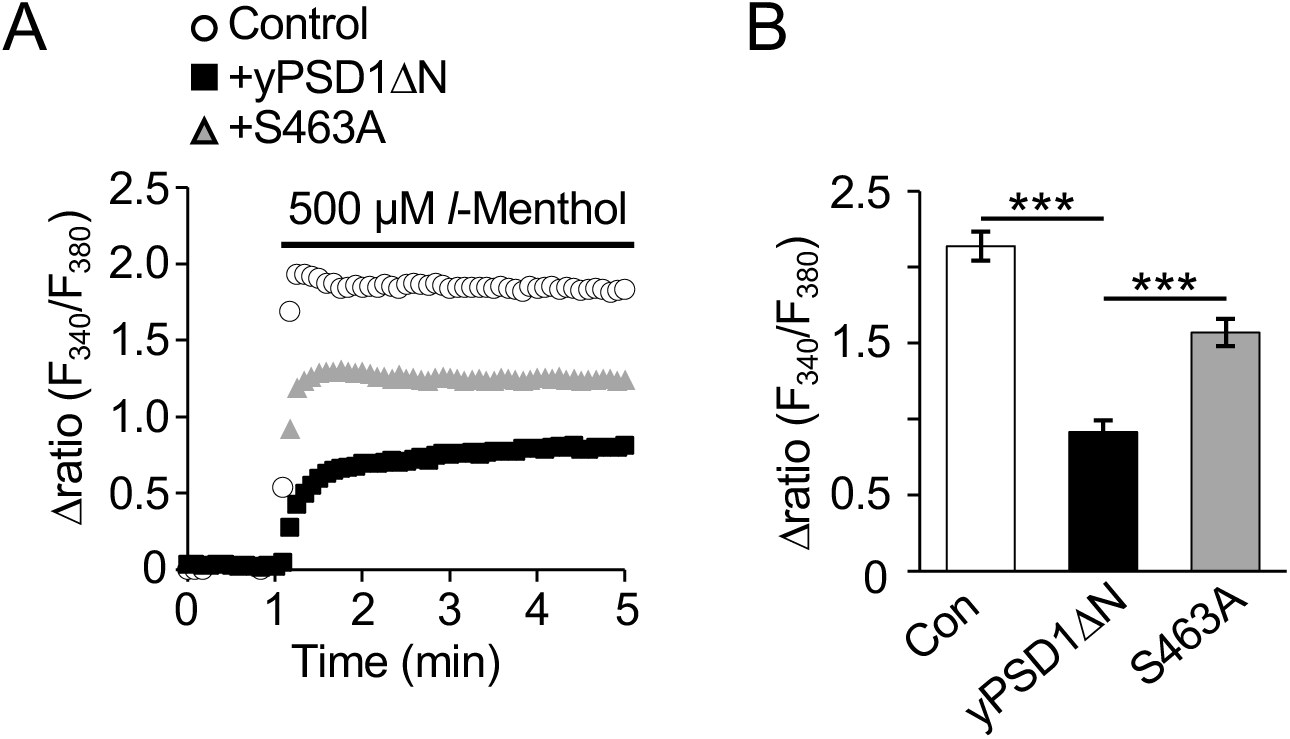
Role of cytosolic PS abundance in the regulation of TRPM8 activity. (A) Fura-2 ratiometric measurements (F340/F380) of *l*-menthol-induced Ca^2+^ mobilization in cells transiently expressing TRPM8 alone (open circles), coexpressing yPSD1ΔN (filled squares), or coexpressing yPSD1ΔN S463A (gray triangles). Average Δratio traces are shown. (B) Maximum Δratio induced by *l*-menthol calculated from (A). Control, n = 79; yPSD1ΔN, n = 58; S463A, n = 73 cells. Mean ± SEM. *** *P* < 0.001.

### Cytoplasmic leaflet of PS regulates the biochemical state of TRPM8

Finally, we investigated the possibility that phospholipid asymmetry regulates the biochemical state of TRPM8. Immunoblot analysis of Myc-tagged TRPM8 detected higher- and lower-molecular-weight bands that were absent from control cells (Fig. S7). When yPSD1ΔN was coexpressed, the intensity ratio of the upper band to the total TRPM8 signal was significantly reduced, whereas coexpression of yPSD1ΔN S463A did not significantly alter this ratio (Fig. 5A, B). Similarly, coexpression of CA-TMEM16F decreased this ratio, whereas wild-type TMEM16F had no significant effect (Fig. 5C, D). Together, these findings suggest that PS in the cytoplasmic leaflet maintains the biochemical state of TRPM8 required for its activity.

**Figure 5.**
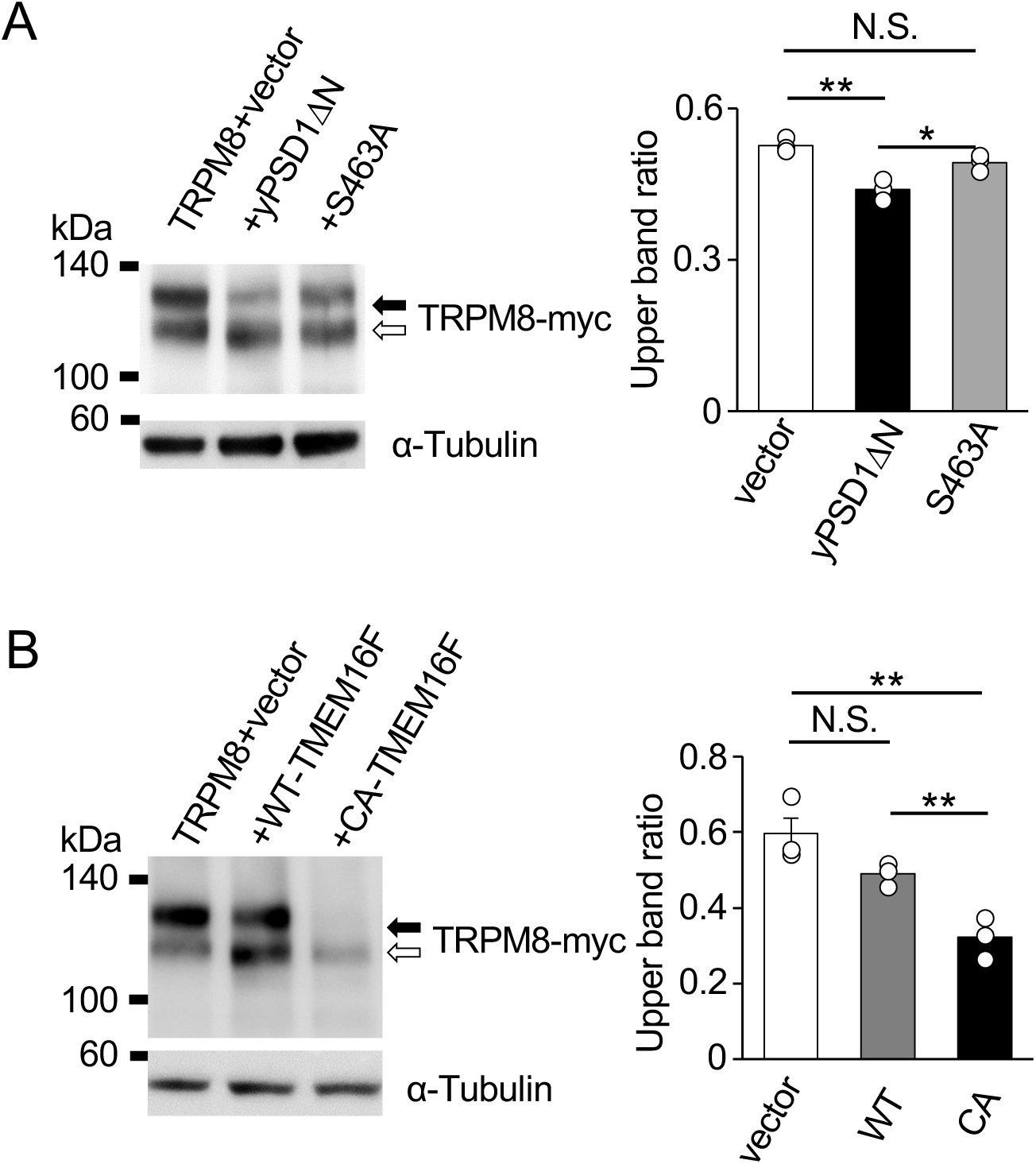
Biochemical state of TRPM8 in phospholipid asymmetry-disrupted cells. Immunoblot analysis of Myc-tagged TRPM8 in cells expressing yPSD1ΔN (A) or TMEM16F (B). The ratio of the upper TRPM8 band to the total TRPM8 signal was quantified in the right panels. The higher- and lower-molecular-weight bands are indicated by black and white arrows, respectively. α-Tubulin was used as a loading control. Mean ± SEM. \**P* < 0.05, \*\**P* < 0.01; N.S., not significant.

## Discussion

Mammalian cells actively maintain phospholipid asymmetry across the cell membrane, implying that this membrane property serves important physiological functions (32). Nevertheless, the molecular targets regulated by phospholipid asymmetry have not been systematically identified. Building on our previous work, we established a screening strategy based on artificial disruption of phospholipid asymmetry and, using this approach, identified TRPM8 as a potential target of regulation by phospholipid asymmetry. Electrophysiological analyses confirmed that disrupting phospholipid asymmetry suppresses agonist-evoked TRPM8 currents (Fig. 2). We further showed that phospholipid asymmetry is required for efficient TRPM8 activation by innocuous cold (Fig. 3). In addition, our results suggest that PS in the cytoplasmic leaflet underlies this requirement for efficient TRPM8 activation (Fig. 4). Finally, biochemical analyses suggested that phospholipid asymmetry maintains the biochemical state of TRPM8 required for efficient channel activation (Fig. 5). Collectively, these findings identify TRPM8 as a previously unrecognized phospholipid asymmetry-regulated ion channel. More broadly, they provide experimental evidence that perturbation of phospholipid asymmetry alters the biochemical state of membrane proteins, thereby expanding our understanding of how phospholipid asymmetry regulates membrane protein function.

In the present study, phospholipid asymmetry was disrupted in HEK293T cells by ectopic expression of the constitutively active phospholipid scramblase CA-TMEM16F, thereby establishing a screening strategy for identifying regulated membrane proteins. A major advantage of this approach is its simplicity. Because only coexpression of CA-TMEM16F and the protein of interest is required, the assay should, in principle, be readily applicable to diverse cell types without relying on endogenous expression of phospholipid transporters, including phospholipid flippases. An important consideration, however, is whether expression of CA-TMEM16F indirectly alters the membrane lipid environment through changes in lipids other than phosphatidylserine. To address this possibility, we examined cytoplasmic PIP₂ using a genetically encoded fluorescent probe (33,34) and detected no measurable change in its abundance (Fig. S8). Consistent with this finding, agonist-induced activation of the PIP₂-regulated channels TRPV1 (35,36), TRPA1 (37) and TRPV4 (38) was unaffected, whereas TRPM8 activity was selectively suppressed (Fig. 1). Although these observations do not exclude subtle alterations in PIP₂ homeostasis, they suggest that the screening strategy preferentially identifies membrane proteins regulated by phospholipid asymmetry rather than by PIP₂-dependent mechanisms. Another limitation is that the approach is restricted to proteins for which robust functional assays are available. To achieve a more comprehensive identification of membrane proteins regulated by phospholipid asymmetry, complementary strategies that overcome these limitations will be required.

Both expression of a constitutively active scramblase and expression of a cytosolically targeted PS decarboxylase suppressed TRPM8-dependent Ca²⁺ influx. These findings prompted us to investigate how disruption of phospholipid asymmetry reduces TRPM8 activity (39). Despite disruption of phospholipid asymmetry, TRPM8 remained localized to the plasma membrane in cells expressing CA-TMEM16F, similar to its localization in control cells, suggesting that its subcellular localization is largely unaffected (Fig. S9). In contrast, immunoblot analysis detected distinct higher- and lower-molecular-weight forms of TRPM8 and showed that disruption of phospholipid asymmetry reduced the abundance of the higher-molecular-weight form (Fig. 5). Because TRPM8 has been reported to undergo N-glycosylation and mutation of its N-glycosylation site reduces channel activity, these observations raise the possibility that disruption of phospholipid asymmetry impairs TRPM8 glycosylation, thereby reducing channel activity. Alternatively, particularly in cells expressing CA-TMEM16F, phospholipid asymmetry disruption may destabilize TRPM8, although this possibility could also reflect an artifact associated with ectopic TRPM8 expression (Fig. 5C). Distinguishing between these possibilities will require future studies using cells expressing endogenous TRPM8 to define more precisely how phospholipid asymmetry regulates TRPM8.

Our findings also provide insight into the feature of phospholipid asymmetry that is critical for TRPM8 regulation. Expression of yPSD1ΔN, which reduces PS abundance in the cytoplasmic leaflet, significantly suppressed TRPM8-dependent Ca²⁺ influx. In contrast, extracellular application of lysophosphatidylserine (lysoPS) had no effect on agonist-induced TRPM8 activation (Fig. S10). Together, these observations suggest that TRPM8 activity depends primarily on maintaining a sufficient concentration of PS within the cytoplasmic leaflet rather than on its distribution between the outer and inner leaflets. Because PS is a relatively minor phospholipid relative to PC and PE, maintenance of phospholipid asymmetry may, at least in part, preserve the threshold level of cytoplasmic PS required for the function of specific membrane proteins such as TRPM8.

Because the present study relied on ectopic expression of TRPM8 in HEK293T cells, it does not directly address the physiological significance of phospholipid asymmetry-dependent regulation of TRPM8. Nevertheless, TRPM8 has been implicated in diverse physiological and pathological processes, including cold sensation (27), neuropathic pain (40), and overactive bladder (41). In addition, our results demonstrate that phospholipid asymmetry selectively regulates TRPM8 among the TRP channels examined (Fig. 1) and is required for activation by both chemical agonists and cold (Fig. 2, 3). Together, these findings support the physiological relevance of our observations. Future studies examining PS dynamics in more physiologically relevant models in which TRPM8 plays particularly important roles will therefore be valuable. More broadly, our results demonstrate the utility of this experimental platform as a screening strategy for identifying membrane proteins regulated by phospholipid asymmetry. Applying this approach to a broader range of membrane proteins should facilitate experimental testing of the flip-flop switch hypothesis and further advance our understanding of the physiological functions of phospholipid asymmetry.

### Experimental procedures Materials

*l*-Menthol, capsaicin, and GSK1016790A were purchased from FUJIFILM Wako Pure Chemical Corporation. Allyl isothiocyanate was purchased from Sigma-Aldrich. LysoPS (18:1) was purchased from Avanti Polar Lipids. All other reagents, chemicals, and antibodies used in this study are listed in Table S1.

### Plasmids

Expression vectors encoding the proteins of interest were either purchased or generated by subcloning the corresponding cDNAs into appropriate expression vectors using restriction enzyme-based or In-Fusion seamless cloning. Plasmids used in this study are listed in Table S2.

### Cell culture

HEK293T cells were maintained in DMEM supplemented with 10% heat-inactivated fetal bovine serum (FBS), 1% GlutaMAX, and 0.5% Penicillin-Streptomycin. Cells were cultured at 37°C in a humidified incubator containing 5% CO₂. Plasmids were transfected using Lipofectamine (Thermo Fisher Scientific). For patch-clamp analysis, cells were co-transfected overnight with pIRES2-ZsGreen1-rat TRPM8 together with pcDNA4 encoding WT, CA, or R499A MmTMEM16F, or with an empty vector, at a plasmid ratio of 1:2 (e.g., 0.5 and 1.0 µg, respectively).

### Electrophysiology

Cells grown on poly-L-lysine-coated coverslips were first mounted in an open chamber (Warner Instruments) and superfused with bath solution containing 140 mM NaCl, 5 mM KCl, 2 mM CaCl₂, 2 mM MgCl₂, 10 mM glucose, and 10 mM HEPES (pH 7.4, adjusted with NaOH). Whole-cell voltage-clamp recordings were performed as described previously (42). The internal solution contained 140 mM KCl, 5 mM EGTA, and 10 mM HEPES (pH 7.4, adjusted with KOH). The membrane potential was held at −60 mV, and voltage ramp-pulses from −100 to +100 mV (300 ms duration) were applied every 3 s. Currents were recorded in the whole-cell configuration using an Axopatch 200B amplifier (Molecular Devices), filtered at 5 kHz, digitized using a Digidata 1440A (Molecular Devices), and acquired with pCLAMP 10 (Axon Instruments). For cold stimulation, the bath temperature was first raised above 30°C by perfusion with warmed solution and subsequently lowered by perfusion with ice-chilled solution. Temperature was monitored with a thermocouple (TA-29; Warner Instruments) positioned within 100 μm of the recorded cell.

### Ca^2+^ imaging

Cells were seeded onto poly-L-lysine-coated coverslips, loaded with Fura-2 AM (5 μM) in DMEM containing 10% FBS for 40 min at 37°C, and then washed with HEPES-buffered saline (HBS). The cells were mounted in an open chamber and superfused with a bath solution containing 140 mM NaCl, 5 mM KCl, 2 mM CaCl₂, 2 mM MgCl₂, 10 mM glucose, and 10 mM HEPES (pH 7.4, adjusted with NaOH). Ratiometric Fura-2 fluorescence imaging (F340/F380) was performed using an inverted microscope (Eclipse Ti2-U; Nikon). Images were acquired every 3 s and analyzed using ImageJ Fiji (version 1.53t). Ca²⁺ influx was quantified as the difference (Δratio) between the Fura-2 ratio at each time point and the mean Fura-2 ratio during the first 1 min of imaging. For the experiments shown in Figs. S3 and S10, Ca²⁺ imaging was performed as described previously (7). For cold stimulation experiments, temperature changes were applied as described for the electrophysiological recordings described above.

### Immunoblot analysis

Cells were washed with PBS and lysed on ice for 60 min in lysis buffer containing 10 mM Tris-HCl (pH 7.4), 1% Triton X-100, 0.1% sodium dodecyl sulfate (SDS), 1% sodium deoxycholate, and 1× cOmplete EDTA-free protease inhibitor cocktail (Nacalai Tesque). The lysates were then centrifuged at 14,000 × *g* for 10 min at 4°C. Subsequently, 40 μl of the supernatant was mixed with 10 μl of SDS sampling buffer 1 (50 mM Tris-HCl (pH 8.0), 50% sucrose, 1% SDS, 5 mM EDTA, 0.4% bromophenol blue), followed by addition of 50 μl of SDS sampling buffer 2 (10 mM Tris-HCl (pH 8.0), 10% sucrose, 0.2% SDS, 1 mM EDTA, 0.08% bromophenol blue, and 60% urea). Samples were separated on a 5%–20% gradient SDS-polyacrylamide gel and subsequently transferred onto polyvinylidene fluoride (PVDF) membranes using Trans-Blot SD Semi-Dry Electrophoretic Transfer Cells (Bio-Rad). Immunoblotting was performed using mouse anti-c-Myc antibody (1:1,500) and rabbit anti-α-tubulin antibody (1:5,000). Bound antibodies were detected using horseradish peroxidase-conjugated anti-rabbit immunoglobulin (Ig) G antibody, anti-mouse IgG antibody, or anti-rat IgG together with SuperSignal West Pico (Thermo Scientific) and Ez-Capture MG (Atto). Band intensities were quantified using ImageJ Fiji (version 1.53t).

### Annexin V labeling

Cells were fixed with 4% paraformaldehyde (PFA) for 10 min at room temperature (RT) and then washed with ice-cold Annexin V-binding (AB) buffer containing 140 mM NaCl, 25 mM CaCl₂, and 10 mM HEPES (pH 7.4, adjusted with NaOH). Cells were subsequently incubated with Annexin-V-EGFP (1:100 in AB buffer) on ice for 30 min. Fluorescence images were acquired using a confocal microscope (LSM 800; Carl Zeiss) and processed with ZEN 2.3 (Carl Zeiss).

### Immunofluorescent analysis

Cells were fixed with 4% PFA in PBS for 10 min at RT and permeabilized with 0.1% Triton X-100 in PBS for an additional 10 min at RT. For PIP₂ labeling, cells were permeabilized by rapid freezing in liquid nitrogen instead of Triton X-100 treatment. Cells were then stained with anti-Myc (1:500), anti-RFP (1:500), and anti-FLAG antibodies (1:500), followed by imaging with a confocal microscope (LSM 800; Carl Zeiss). Fluorescence images were processed using ZEN 2.3 (Carl Zeiss).

### Statistical analyses

All statistical analyses were performed using GraphPad Prism 5 (GraphPad Software). Statistical significance between group means was evaluated using a two-sided unpaired *t*-test. For multiple comparisons, analysis of variance (ANOVA) followed by Tukey’s test was performed. *P* values of \**P* < 0.05, \*\**P* < 0.01, and \*\*\**P* < 0.001 were considered statistically significant.

### Data availability

The data that support the findings of this study are available from the corresponding author upon reasonable request.

### Supporting information

This article contains the following supporting information; Figures S1-10 and Tables S1-2.

## Acknowledgements

We thank members of the Hara laboratory for their scientific contributions. This study was supported by the Grant-in-Aid for Scientific Research KAKENHI (22H03484, 25K02994); Grant-in-Aid for Transformative Research Areas

B (23H03854, 23H03856); Intramural Research Grant (8–13) for Neurological and Psychiatric Disorder of NCNP; grants from Takeda Science Foundation, ONO Medical Research Foundation, Uehara memorial foundation, Chugai Foundation for Innovative Drug Discovery Science (SRG2022), and the Asahi Glass Foundation (to Y.H.); and the Grant-in-Aid for Scientific Research KAKENHI (22K20636, 23K14184, 25K18466); grants from UBE Foundation, ONO Medical Research Foundation, Takeda Science Foundation (to A.M.).

## Author contributions

Conceptualization: M.U., Y.H.; Methodology: M.T., M.S., A.S., K.N., T.T., and K.U.; Investigation: R.N., A.M., and E.S.; Supervision: A.M., Y.H.; Writing—original draft: A.M., R.N.; Writing—review and editing: A.M., R.N., and Y.H.

## Abbreviations

TRP: (transient receptor potential)
M8: (Melastatin 8)
PS: (phosphatidylserine)
PC: (phosphatidylcholine)
PE: (phosphatidylethanolamine)
PI: (phosphatidylinositol)
ATP: (adenosine triphosphate)
V2: (Vanilloid 2)
C5: (Canonical 5)
PIP_2_: (phosphatidylinositol 4,5-bisphosphate)
V1: (Vanilloid 1)
V5: (Vanilloid 5)
ML1: (Mucolipin 1)
TMEM16F: (transmembrane protein 16F)
WT: (wild-type)
CA: (constitutively active)
AITC: (allyl isothiocyanate)
A1: (Ankyrin 1)
XKR: (XK-related protein)
Aa: (*Aedes albopictus*)
lysoPS: (lysophosphatidylserine)
FBS: (fetal bovine serum)
SDS: (sodium dodecyl sulfate)
PVDF: (polyvinylidene fluoride)
PFA: (paraformaldehyde)
AB: (Annexin V-binding)

