## Supporting information for "Identification of a transient receptor potential channel that is regulated by phospholipid asymmetry"

### Nakanishi et al. Figure. S1

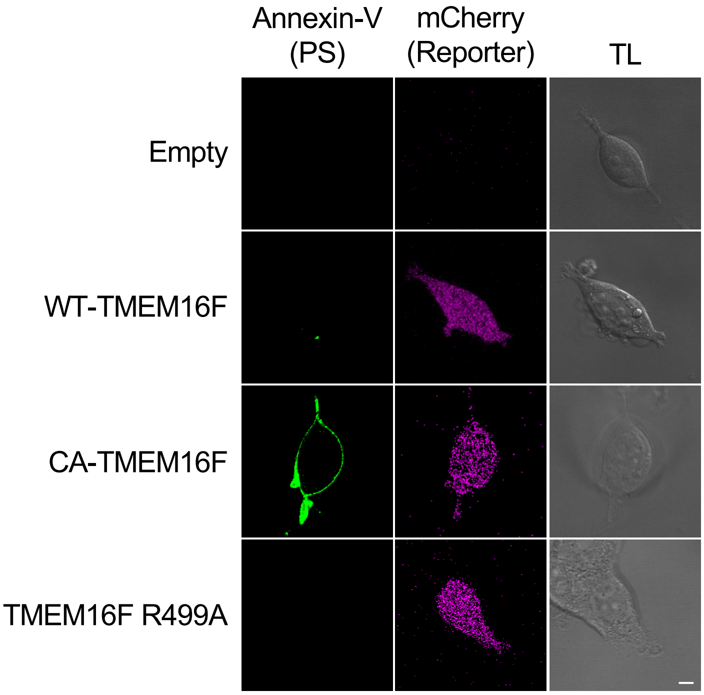

**Figure S1. PS exposure in cells expressing phospholipid scramblases.**

Detection of PS on the extracellular leaflet of the plasma membrane using Annexin V-EGFP. TMEM16F constructs containing an internal ribosome entry site (IRES) followed by mCherry were used to identify the indicated TMEM16F-expressing cells. WT, wild-type; CA, constitutively active; R499A, phospholipid scrambling-deficient mutant. Bars: 5 μm.

Nakanishi et al. Figure. S2

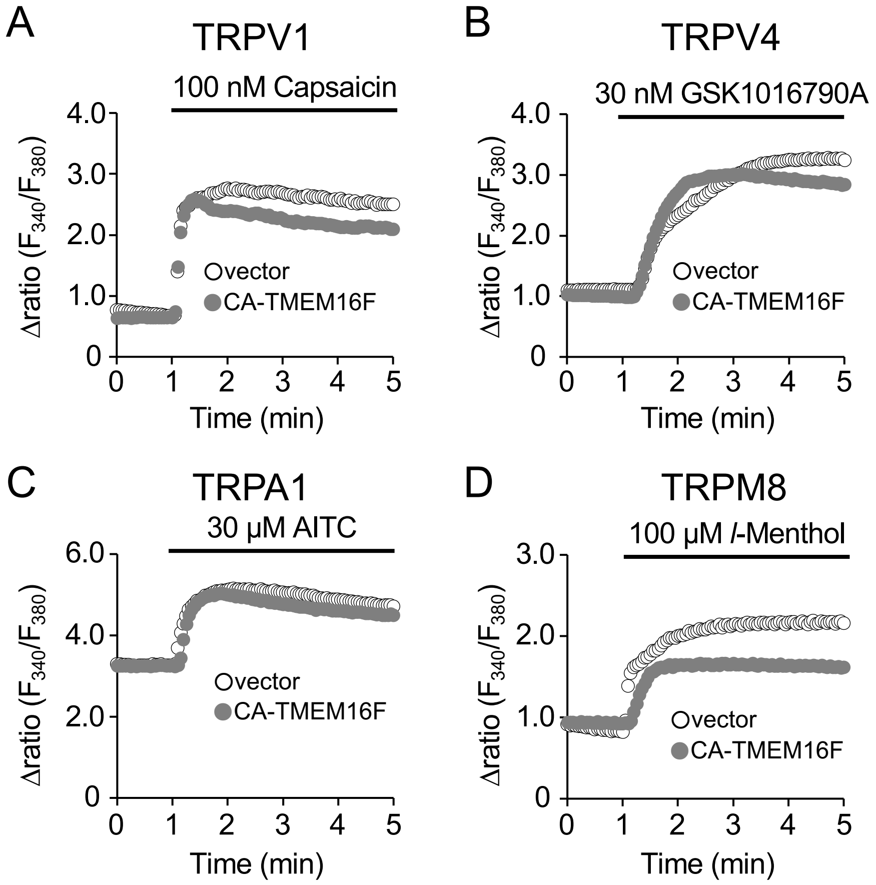

**Figure S2. Representative $Ca^{2+}$ imaging traces used for the TRP channel screen.**

Fura-2 ratiometric measurements (F340/F380) of agonist-induced $Ca^{2+}$ mobilization in cells transiently expressing the indicated TRP channels alone (open circles) or coexpressing CA-TMEM16F (gray circles). Data are presented as the mean ± SEM. These data were used to calculate the Fura-2 Δratio shown in Figs. 1C–F. CA, constitutively active.

Nakanishi et al. Figure. S3

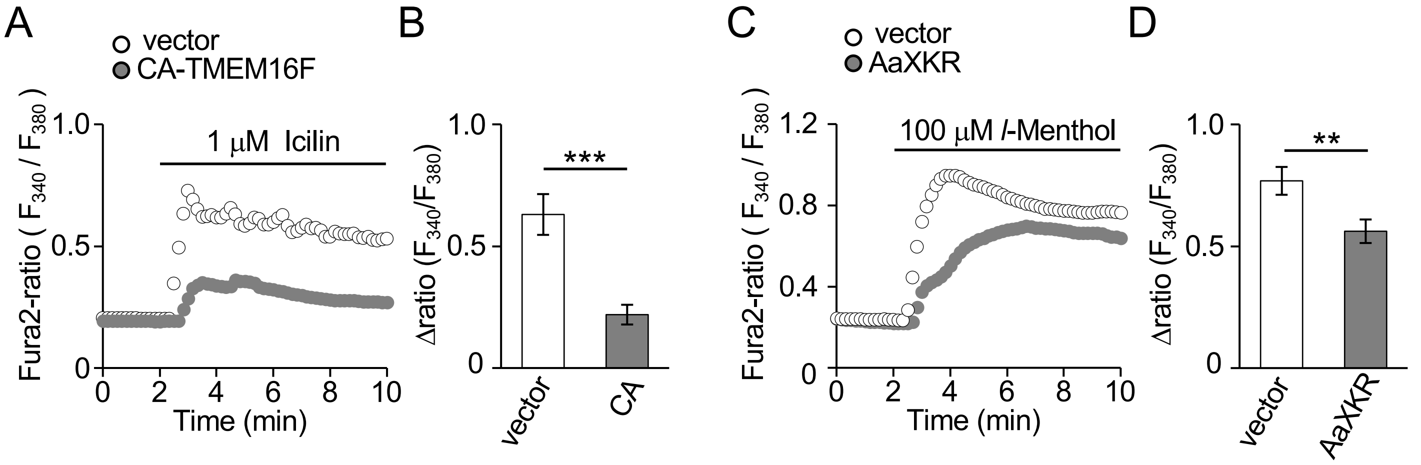

**Figure S3. Suppression of TRPM8-mediated $Ca^{2+}$ influx in phospholipid asymmetry-disrupted cells.**

(A) Fura-2 ratiometric measurements (F340/F380) of icilin-induced $Ca^{2+}$ mobilization in cells transiently expressing TRPM8 alone (open circles) or coexpressing CA-TMEM16F (gray circles). The corresponding Fura-2 Δratio is shown in (B). vector, n = 41; CA-TMEM16F, n = 56 cells. (C) Fura-2 ratiometric measurements (F340/F380) of *l*-menthol-induced $Ca^{2+}$ mobilization in cells transiently expressing TRPM8 alone (open circles) or coexpressing AaXKR (gray circles). The corresponding Fura-2 Δratio is shown in (D). vector, n = 24; AaXKR, n = 39 cells. Mean ± SEM. $^{**}P < 0.01$, $^{***}P < 0.001$. CA, constitutively active. Aa, *Aedes albopictus*.

### Nakanishi et al. Figure. S4

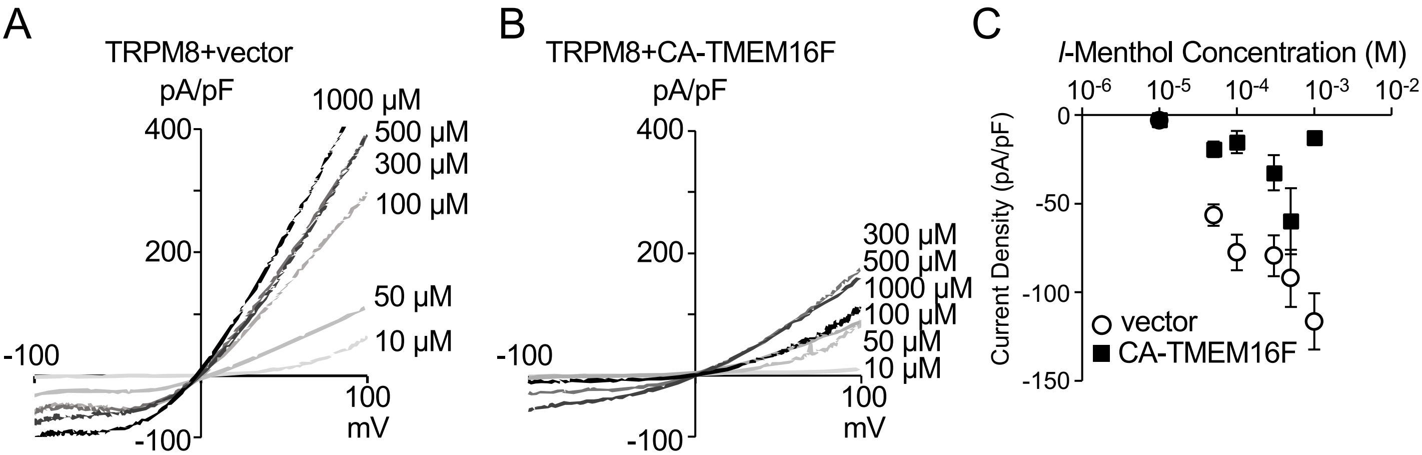

**Figure S4. Dose-dependence of TRPM8 currents evoked by *l*-menthol.**

(A, B) Representative I-V curves of TRPM8 currents induced by *l*-menthol at the indicated concentrations in cells transiently expressing TRPM8 alone (A) or coexpressing CA-TMEM16F (B). The membrane potential was held at −60 mV. (C) Current density of TRPM8 induced by *l*-menthol at the indicated concentrations in cells transiently expressing TRPM8 alone (open circles) or coexpressing CA-TMEM16F (filled squares). Mean ± SEM (n = 4-7). CA, constitutively active.

Nakanishi et al. Figure. S5

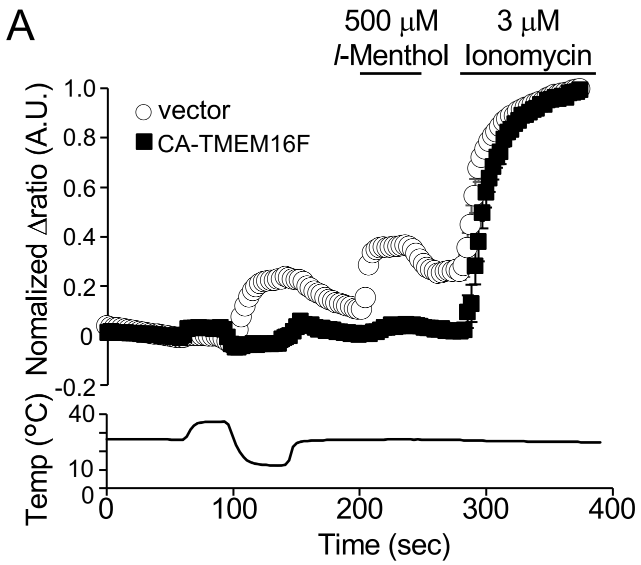

**Figure S5. Cold-induced mouse TRPM8 activation in phospholipid asymmetry-disrupted cells.**

(A) Fura-2 ratiometric measurements (F340/F380) of $Ca^{2+}$ mobilization in cells transiently expressing TRPM8 alone (open circles) or coexpressing CA-TMEM16F (filled squares). Cells were sequentially exposed to cold stimulation, *l*-menthol, and ionomycin. The Fura-2 ratio was normalized to the value obtained following ionomycin application. Average Δratio traces are shown. The maximum Fura-2 Δratio during cold stimulation is shown in (B). vector, N = 6; CA-TMEM16F, N = 5. Mean ± SEM. ****P* < 0.001. CA, constitutively active.

Nakanishi et al. Figure. S6

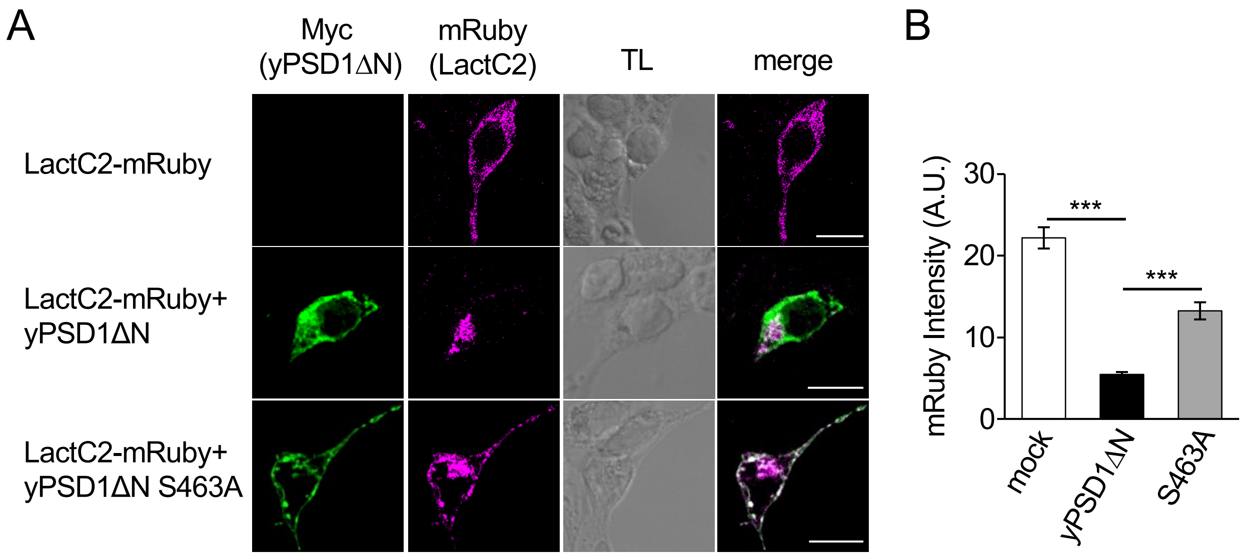

**Figure S6. Effect of yPSD1ΔN expression on cytosolic PS abundance.**

(A) Detection of cytosolic PS using the PS-binding domain of lactadherin fused to mRuby (LactC2; magenta). mRuby fluorescence intensities were shown in (B). yPSD1ΔN S463A represents the catalytically inactive mutant of yPSD1ΔN. mock, n = 86; yPSD1ΔN, n = 115; S463A, n = 129 cells. Mean ± SEM. $**P < 0.01$, $***P < 0.001$. Bars: 10 μm.

Nakanishi et al. Figure. S7

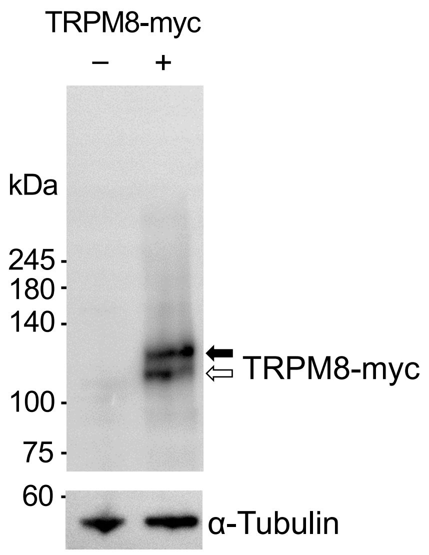

**Figure S7. Biochemical state of ectopically expressed TRPM8.**

Immunoblot analysis of rat TRPM8 expressed in HEK293T cells. TRPM8 protein was detected using an anti-Myc antibody. The higher- and lower-molecular-weight bands detected specifically in TRPM8-expressing cells are indicated by black and white arrows, respectively. α-Tubulin was used as a loading control.

Nakanishi et al. Figure. S8

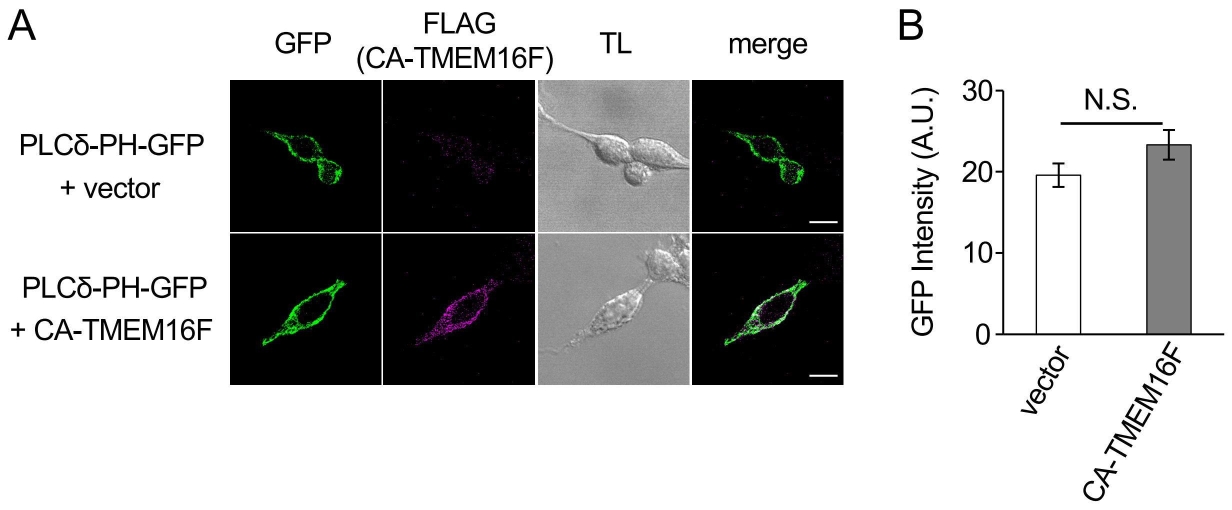

**Figure S8. Cytosolic PI(4,5)$P_2$ levels in cells expressing CA-TMEM16F.**

(A) Detection of cytosolic PI(4,5)$P_2$ using the PH domain of PLCδ (PLCδ-PH; green) in cells with or without expression of FLAG-tagged CA-TMEM16F (magenta). GFP fluorescence intensities were shown in (B). Bars: 10 μm. n = 60 cells per group. Mean ± SEM. N.S.., not significant. CA, constitutively active.

### Nakanishi et al. Figure. S9

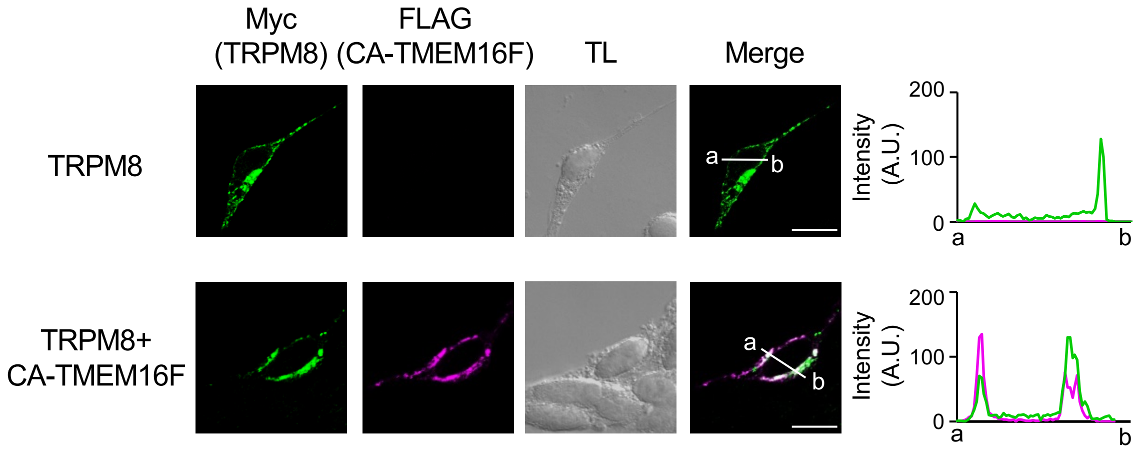

**Figure S9. Localization of TRPM8 in cells expressing CA-TMEM16F.**

Detection of transiently expressed Myc-tagged TRPM8 (green) in cells with or without expression of FLAG-tagged CA-TMEM16F (magenta). Fluorescence intensity profiles of Myc and FLAG along the line indicated by a-b are shown in right panels. Bars: 10 µm. CA, constitutively active.

Nakanishi et al. Figure. S10

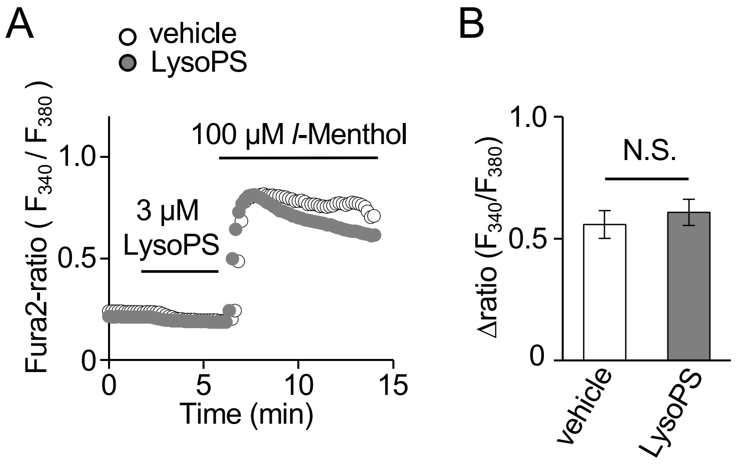

**Figure S10. Effect of extracellular lysoPS application on TRPM8-mediated $Ca^{2+}$ influx.**

(A) Fura-2 ratiometric measurements (F340/F380) of *l*-menthol-induced $Ca^{2+}$ mobilization in cells transiently expressing TRPM8. Before *l*-menthol stimulation, cells were pretreated with vehicle (open circles) or 3 μM lysoPS (gray circles). The corresponding Fura-2 Δratio is shown in (B). vehicle, n = 32; LysoPS, n = 27 cells. Mean ± SEM. N.S.., not significant.

Table S1. Plasmids used in this study.

| Plasmid | Source | Insert | | Vector |
|---|---|---|---|---|
| pIRES2-ZsGreen1-RatTrpm8 | Handmade | Addgene #64879 | | Clontech 632478 |
| pIRES2-mCherry | Handmade | Addgene #49243 | | Clontech 632478 |
| pIRES2-mCherry-RatTrpm8 | Handmade | Addgene #64879 | | Clontech 632478 |
| pIRES2-mCherry-RatTrpv1 | Handmade | Addgene #79649 | | pIRES2-mCherry |
| pIRES2-mCherry-HsTRPV4 | Handmade | cDNA | | pIRES2-mCherry |
| pIRES2-mCherry-HsTRPA1 | Handmade | cDNA | | pIRES2-mCherry |
| pcDNA4-myc-His-A | invitrogen #V863-20 | | | |
| pcDNA4-MmTMEM16F-CA-FLAG | Handmade | pMXs-puro-MmTMEM16F-CA-FLAG | Tsuchiya, T. et al. Nat Commun 2018 | pcDNA4-Myc-His-A |
| pcDNA4-MmTMEM16F-FLAG | Handmade | pcDNA4-MmTMEM16F-CA-FLAG | | pcDNA4-Myc-His-A |
| pcDNA4-MmTMEM16F-CA-FLAG (R499A) | Handmade | pcDNA4-MmTMEM16F-CA-FLAG | | pcDNA4-Myc-His-A |
| pcDNA5/FRT-MmTrpm8 | Handmade | cDNA | | pcDNA5/FRT |
| pHD-DsRed | Addgene #13771 | | | |
| pcDNA3-Myc-GGGSx3-yPSD1 (aa99-500) | Matsudaira, T. et al. Nat Commun (2017) | | | |
| pcDNA3-Myc-GGGSx3-yPSD1 (aa99-500) (S463A) | Matsudaira, T. et al. Nat Commun (2017) | | | |
| pcDAN4-IRES-mCherry | Handmade | pIRES2-mCherry | | pcDNA4-Myc-His-A |
| pcDAN4-MmTMEM16F-FLAG-IRES-mCherry | Handmade | pcDNA4-MmTMEM16F-FLAG | | pcDAN4-IRES-mCherry |
| pcDAN4-MmTMEM16F-CA-FLAG-IRES-mCherry | Handmade | pcDNA4-MmTMEM16F-CA-FLAG | | pcDAN4-IRES-mCherry |
| pcDAN4-MmTMEM16F-CA-FLAG (R499A)-IRES-mCherry | Handmade | pcDNA4-MmTMEM16F-CA-FLAG (R499A) | | pcDAN4-IRES-mCherry |
| pcDAN4-AaXKR | Handmade | pIRES2-puro3-EGFP-AaXKR | Shiomi, A. et al. Cel Rep 2021 | pcDNA4-Myc-His-A |
| pmRuby2-C1-LactC2 | Addgene #22852 | | | |
| pClover-N1-PHhsplcd1 | Addgene #22852 | | | |

Table S2. Materials used in this study.

| REAGENT or RESOURCE | SOURCE | #IDENTIFIER |
|---|---|---|
| **Antibodies** | | |
| Polyclonal anti-Myc-tag | MBL | #562 |
| Monoclonal anti-c-myc clone 9E10 | Sigma-Aldrich | #M5546 |
| Monoclonal anti-RFP | MBL | #M208-3 |
| Monoclonal anti-FLAG® M2 | Sigma-Aldrich | #F1804-200UG |
| Plyclonal anti-α-Tubulin | CST | #2144 |
| Anti-rabbit IgG, HRP-linked antibody | CST | #7074 |
| Anti-mouse IgG, HRP-linked antibody | CST | #7076 |
| Anti-mouse IgG, Alexa Fluor 488 | Thermo Fisher Scientific | #A21424 |
| Anti-rabbit IgG, Alexa Fluor 555 | Thermo Fisher Scientific | #A11008 |
| **Chemicals** | | |
| *l*-Menthol | FUJIFILM | #132-03752 |
| Capsaicin | FUJIFILM | #030-11353 |
| GSK1016790A | FUJIFILM | #073-06491 |
| Allyl isothiocyanate | Sigma-Aldrich | #377430-100G |
| LysoPS (18:1) | Avanti | #858143P |
| Ionomycin | FUJIFILM Wako | #095-05831 |
| Sodium Chloride | FUJIFILM Wako | #191-01665 |
| Potassium Chloride | FUJIFILM Wako | #163-03545 |
| Calcium Chloride | FUJIFILM Wako | #038-24985 |
| Calcium Chloride Dihydrate | FUJIFILM Wako | #033-25035 |
| Magnesium Chloride Hexahydrate | FUJIFILM Wako | #135-00165 |
| Sodium Hydroxide | FUJIFILM Wako | #198-13765 |
| D(+)-Glucose | FUJIFILM Wako | #049-31165 |
| GlutaMAX™ Supplement | Gibco | #35050061 |
| 2-[4-(2-Hydroxyethyl)-1-piperazinyl]ethanesulfonic acid | FUJIFILM Wako | #GB70 |
| 2-Amino-2-hydroxymethyl-1,3-propanediol | FUJIFILM Wako | #207-06275 |
| GEDTA(EGTA) | Dojindo | #G002 |
| EDTA 2Na | Dojindo | #N001 |
| Ethanol | FUJIFILM Wako | #057-00456 |
| Methanol | FUJIFILM Wako | #137-01823 |
| ply-L-lysine | Sigma-Aldrich | #P4707 |
| Dimethyl Sulfoxide | FUJIFILM Wako | #045-28335 |
| 4% Paraformaldehyde Phosphate Buffer Solution | FUJIFILM Wako | #163-20145 |
| Triton® X-100 | MP Biomedicals | #04807426-CF |
| Polysorbate 20 | MP Biomedicals | #02103168-CF |
| Sodium Dodecyl Sulfate | FUJIFILM Wako | #192-13981 |
| Sodium Deoxycholate | FUJIFILM Wako | #190-08313 |
| cOmplete, EDTA-free Protease Inhibitor Cocktail | Roche | #11873580001 |
| **Reagents** | | |
| Lipofectamine™ Transfection Reagent | Invitrogen | #18324012 |
| Lipofectamine™ 3000 Transfection Reagent | Invitrogen | #L3000015 |
| PLUS™ Reagent | Invitrogen | #11514015 |
| AnnexinV-EGFP | VioVision | #1004-200 |
| Fura2-AM | DOJINDO | #F015 |
| Fetal bovine serum | Gibco | #A5256701 |
| Bovine Serum Albumin (IgG-Free,Protease-Free) | Jackson ImmunoResearch | #001-000-162 |
| DMEM (High Glucose) | FUJIFILM Wako | #041-30081 |
| DMEM (High Glucose) without L-Glutamine and Phenol Red | FUJIFILM Wako | #040-30095 |
| SuperSignal™ West Pico PLUS Chemiluminescent Substrate | Thermo Fisher Scientific | #34580 |
| Opti-MEM | Gibco | #31985070 |
| D-PBS(-) | FUJIFILM Wako | #049-29793 |
